# Restoration of Fbxo7 expression in dopaminergic neurons restores tyrosine hydroxylase in a mouse model of Parkinson’s disease

**DOI:** 10.1101/2024.09.28.615591

**Authors:** Sara Al Rawi, Pamela Tyers, Roger A Barker, Heike Laman

## Abstract

Mutations in *FBXO7* are linked to an atypical parkinsonism. Conditional knock out (KO) of Fbxo7 in dopaminergic neurons in a mouse model caused a neurodegenerative phenotype, including a significant reduction in striatal TH staining at 6 weeks of age and a significant loss of dopaminergic neurons in the SNpc. To test whether re-expression of Fbxo7 could act as a treatment to prevent or restore TH expression in the striatum in this model, we used a rAAV vector to deliver murine Fbxo7 and a mRuby fluorescent marker to dopaminergic neurons. We found that Fbxo7 expression, both before and after the TH loss, restored its expression in the striatum and nucleus accumbens in the mouse. This study therefore highlights that Fbxo7 is important for the integrity and persistence of the dopaminergic nigrostriatal pathway in the mammalian brain, which could be of relevance to Parkinson’s disease with therapeutic implications.

## Introduction

Most cases of Parkinson’s disease (PD) occur in individuals over the age of 50 with the majority being sporadic. However, approximately 1 in 20 people with Parkinson’s (PwP) is under the age of 40, and their disease often has a genetic cause. These rare genetic forms often target specific cellular systems, which has also been implicated in sporadic PD. These include autophagy, mitochondrial and proteostasis, oxidative stress responses, neuro-inflammation, and intracellular trafficking (1-3). Moreover, these systems are inter-linked such that dysfunction of one, e.g. protein homeostasis, may lead to or exacerbate dysfunction of others, e.g., autophagy and stress responses.

To investigate the pathological mechanisms underlying the loss of dopaminergic (DA) neurons in PD, we developed a murine model with conditional loss of one of the PARK genes, PARK15/Fbxo7 (4). It progressively develops many of the hallmark features of human idiopathic PD patients including an early reduction in TH+ cell size in the substantia nigra pars compacta (SNpC) neurons with loss of fibre innervation of the striatum and associated dopamine levels by 6 weeks. This is then followed by significant DA cell loss in SNpC at 20 weeks. Our *Dat* ^*Cre*^ *Fbxo7* ^−/fl^ mouse shows clear cellular nigrostriatal degeneration, and together with other conditional Fbxo7 KO models in other neuronal cell populations, argues for its critical role in neurons (4-7).

Fbxo7 is a multi-functional protein that impacts on several suspected pathways implicated in the aetiology of PD, ranging from cell cycle regulation to mitophagy to proteostasis (8, 9). Moreover, patients with mutations in Fbxo7/PARK15 present with a wide range of clinical signs and symptoms that, in some cases, are indistinguishable from PD clinically (3, 10-27). This suggests that Fbxo7 plays a critical role both in neuronal development and provides neuroprotection later in life. We therefore sought to address whether using rAAV to restore Fbxo7 expression in our conditional KO mice would prevent the striatal denervation seen at 6 weeks and whether the re-expression of Fbxo7 after the onset of the striatal phenotype would restore its TH expression. We found the treatment of mice with rAAV encoding Fbxo7, both before and after the loss of TH+ staining in the striatum, robustly restored TH expression in the midbrain. These data indicate that Fbxo7-regulated pathways are neuroprotective before and after the onset of an overt phenotype.

As such elucidating the relevant Fbxo7-regulated pathway may open up new therapeutic avenues for PD patients.

## Results

Previously, we reported that Fbxo7 loss in DA neurons leads to abnormal nigrostriatal development manifesting from 6 weeks of age at the level of the striatum (4). Specifically, whereas *Dat*^*Cre*^ *Fbxo7* ^−/+^ mice, hereafter referred to as control mice, show robust and uniform TH expression across the striatum, in *Dat*^*Cre*^ *Fbxo7* ^−/fl^ mice, referred to as conditional KO (cKO) mice, there is significantly less TH expression at 6 weeks of age. By 12 weeks, when striatal TH^+^ fibre density plateaus in control mice, cKO mice showed significantly decreased density of TH^+^ fibres in the striatum and nucleus accumbens. We therefore assessed whether Fbxo7 introduction could prevent the early TH loss in cKO mice. We first verified that there was no decrease in TH+ staining in the striatum of cKO at weaning, i.e., at 3-4 weeks of age, the interval during which we could inject mice. We confirmed that the mean optical intensity used as a measure of TH+ fibre density in the striatum was comparable for cKO and control mice at 3-4-weeks of age (data not shown). We then generated rAAV constructs that allowed Cre-dependent expression from an inverted open reading frame (i.e., ‘flexed’), which expressed mRuby as a fluorescent tracer of expression and murine Fbxo7, by ribosomal skipping induced by a P2A site (see Materials and Methods for details). This construct directs re-expression of mFbxo7 specifically to the DAT-Cre-expressing cells. 4-week-old mice were intravenously injected with an inoculum of AAV9-PHP.eB.CAG.DIO.P2A.mRuby (Empty Vector (EV) virus) or with AAV9-PHP.eB.CAG.DIO.FBXO7.P2A.mRuby (Fbxo7 virus) and left to recover for 6 weeks to allow them to fully express the transgene *in vivo* prior to harvest. To verify expression of the transgenes delivered by the rAAV, brain sections were subjected to immunofluorescence assays using primary antibodies to TH, while mRuby fluorescence was directly visualized. Expression of mRuby was evident and restricted to the midbrains of mice injected with the Fbxo7 virus (Figure 1A). Moreover, this fluorescence co-localised with the TH+ cells in the substantia nigra (Figure 1A, B). To quantify TH+ staining in the striatum, immunohistochemistry was performed on brain sections and visualized by diaminobenzidine (DAB) staining (Figure 1C). Mean optical density across the dorsal striatum and the nucleus accumbens was measured using TH-positive stained sections and used to determine the fibre density (see Materials & Methods). Consistent with previous findings, cKO mouse exhibit a significant 34.5% reduction in TH+ staining in the striatum and 44.4% reduction in the nucleus accumbens at 10 weeks, compared to control mice (Figure 1D, i vs. ii). Treatment of cKO mice by injection with Fbxo7 virus resulted in a 204% and 202% increase in staining in the striatum and nucleus accumbens, respectively, relative to untreated cKO mice (Figure 1D, ii vs. iii). When this was compared to the levels of staining in control mice (Figure 1D, ii vs. iv), this represented a restoration to the amount of staining, of around ∼110% in both the striatum and nucleus accumbens as observed in the control brains. Although the injection of the EV virus caused a slight elevation in the amount TH+ staining in the cKO mouse, this was not statistically significant and TH+ levels were still reduced by 24.7% and 33.8% in the striatum and nucleus accumbens, respectively, compared to control mice (Figure 1D, iii vs. iv). These data show that the expression of Fbxo7 prior to the onset of the loss of striatal TH expression prevented this phenotype from developing out to 6 weeks after treatment.

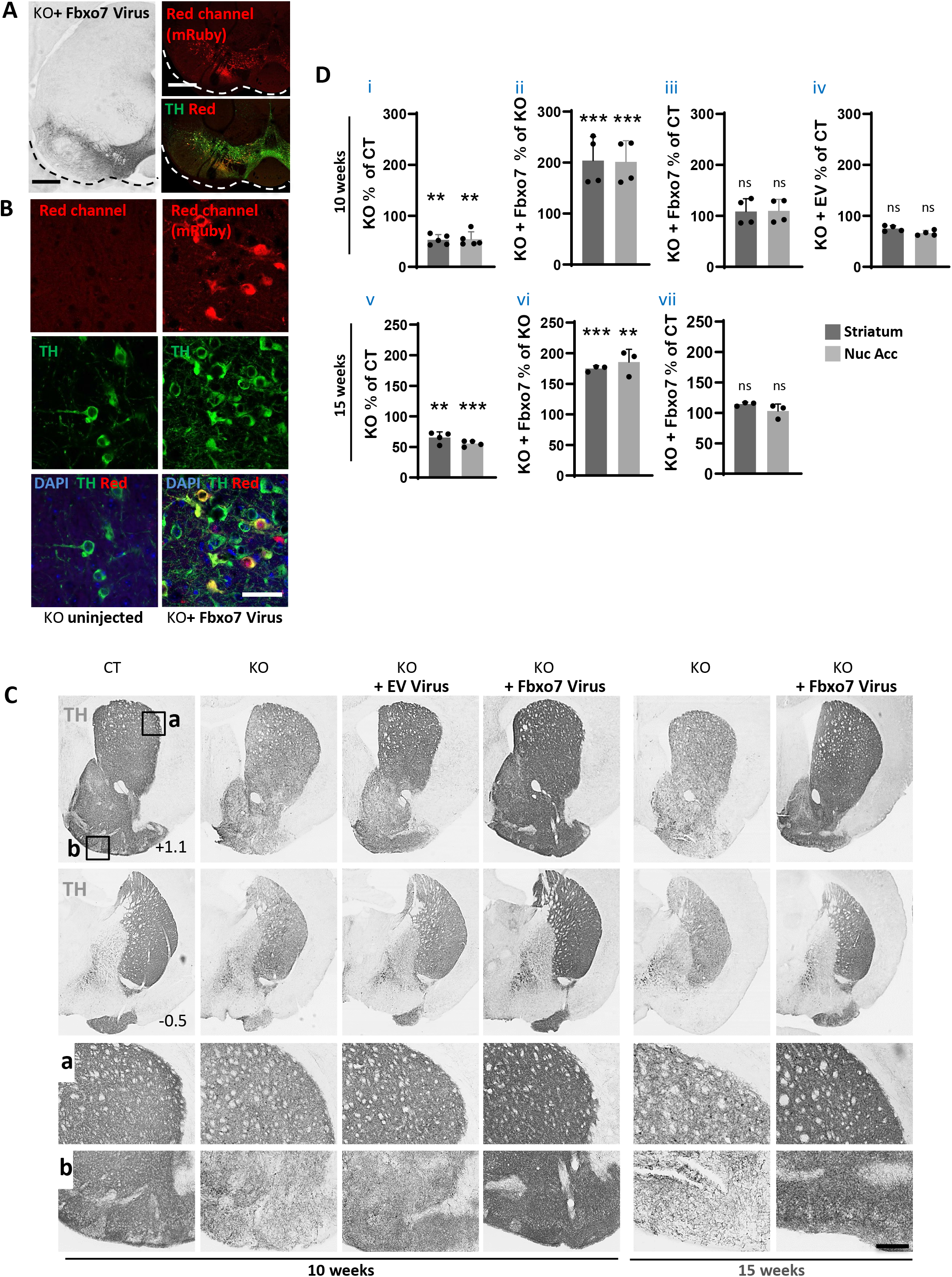
Restoration of Fbxo7 expression rescues TH positive fibre innervation in the striatum of Fbxo7-deficient mice. **(A)** TH immunostaining (DAB, dark grey on the left panel; Alexa Fluor 488, green on the lower right panel) in mouse brain sections at the level of the substantia nigra pars compacta (SNpc), from a mouse injected with the Fbxo7 virus. mRuby expression is observed in red (right panels). **(B)** Magnified view of the TH+ neurons in the SNpc (TH, Alexa Fluor 488, green). No signal in red is observed in the brain section from uninjected KO mice (KO; *Dat*^*Cre*^ *Fbxo7*^*fl/-*^) while mRuby expression is observed in the brain from KO mice (KO; *Dat*^*Cre*^ *Fbxo7*^*fl/-*^) injected with Fbxo7 virus. The scale bar in (A) is 500μm and in (B) is 50μm. **(C)** TH-DAB-immunostained sections of the striatum at +1.1 mm (top panels) and −0.5 mm (bottom panels) relative to the bregma from a control mouse (CT; *Dat*^*Cre*^ *Fbxo*^*+/-*^) at 10 weeks, and conditional knock-out mice (KO; *Dat*^*Cre*^ *Fbxo7*^*fl/-*^) at 10 and 15weeks of age, uninjected or injected with rAAVs, as indicated. (a, b) Magnified regions (boxed in (C)), showing the TH+ fibres in the striatum (a) and olfactory tubercle (b). **(D)** Optical density quantification of TH+ fibre density in KO (*Dat*^*Cre*^ *Fbxo7*^*fl/-*^) mice at 10 or 15 weeks of age, uninjected or injected with rAAVs, as indicated. Results are presented as a percentage of control (*Dat*^*Cre*^ *Fbxo*^*+/-*^) or percentage of KO (*Dat*^*Cre*^ *Fbxo7*^*fl/-*^) at each age with or without rAAV injection, in the striatum and nucleus accumbens (Nuc Acc). The scale bar in (b) represents 800 μm for the two top panels of (C) and 300 μm for a, b. **p < 0.01; ***p < 0.001; ns = not significant.

We next tested whether the re-expression of Fbxo7 using rAAV could act as a therapeutic agent to restore TH+ expression after its significant loss is already apparent in the striatum. cKO mice showed a 30% reduction at 6 weeks and 44% reduction at 12 weeks in TH staining in the striatum (4). We therefore injected the cKO mice at 9 weeks, during an interval when TH expression is declining, and as before, allowed a 6-week recovery period prior to harvesting brains for analysis. At 15 weeks, cKO mice show the expected 34.5% and 44.4% reduction in TH+ staining in the striatum and nucleus accumbens (Figure 1D, v). Remarkably, brains from cKO mice treated with an inoculum of the Fbxo7 virus showed robust TH staining in both regions assessed. The levels of this increase were 174.8% in the striatum and 185% in the nucleus accumbens, representing over a 3-fold increase relative to the TH+ levels in the cKO mouse (Figure 1D, v vs. vi). When compared to control mice, this corresponded to a restoration of TH expression to 114.5% in the striatum and 103.1% in the nucleus accumbens of the levels seen in control mice (Figure 1D, vii). These data demonstrate that the re-expression of Fbxo7 in cKO, in animals which already have a significant reduction in TH in the striatum and nucleus accumbens, can restore its expression in both regions. Our data suggest that Fbxo7-regulated pathways are therapeutically relevant in modulating TH expression in the nigrostriatal pathway even when it is failing.

## Discussion

Fbxo7 is a multifunctional protein with critical physiological roles in a variety of cell types, including lymphocytes, sperm, erythrocytes, and dopaminergic neurons. Whether the same Fbxo7-regulated pathways drive its essential functions across these diverse cell types remains unclear, although there may be unexpected overlaps in the cellular machinery and regulatory processes required for their development. The mechanistic basis of Fbxo7 neuropathology has been linked to its dual impact on mitochondrial homeostasis and proteasome dysfunction (5, 6, 10, 28-32). Unlike PINK1 and Parkin knockout mice—which, despite showing mitochondrial defects like increased damage and impaired oxidative stress response in the ventral midbrain, do not exhibit nigrostriatal degeneration (33-37) —Fbxo7 conditional knockout (cKO) models demonstrate pronounced nigrostriatal and neurodegeneration, more closely mimicking Parkinson’s disease (4-6). This divergence suggests that either robust compensatory mechanisms exist in PINK1 and Parkin KO mice, or that the simultaneous failure of multiple interlinked neuroprotective pathways, as seen with Fbxo7 dysfunction, is required to drive neurodegeneration. The dual disruption of mitochondrial and proteasomal systems by Fbxo7 may explain the unique neurodegenerative outcomes observed in these models (4-6).

In our study, we used the DAT^Cre^ Fbxo7 cKO mouse, which exhibits progressive nigrostriatal degeneration, to investigate whether reintroducing Fbxo7—either before or after the onset of nigrostriatal degeneration—could prevent or reverse these defects. Remarkably, the use of recombinant AAV (rAAV) to deliver Fbxo7 to dopaminergic neurons both prevented the loss of TH expression in the striatum and restored TH levels in cases where TH loss had already occurred. These findings provide compelling evidence that restoring Fbxo7-regulated pathways can exert a therapeutic effect in a progressive neurodegenerative condition. This underscores the importance of identifying the specific Fbxo7-regulated pathway(s) that are neuroprotective, with future experiments aimed at testing therapeutic interventions in these pathways to mitigate PD-associated phenotypes, including neuronal loss and behavioral deficits in Fbxo7 models of parkinsonism.

## Materials and Methods

### Mice

All experiments with mice were performed in accordance with the UK Animals (Scientific Procedures) Act 1986 and ARRIVE guidelines. Mice were housed in individually ventilated cages with unrestricted access to food and water, and a 12-hour day-night cycle. Animal licenses were approved by the Home Office and the University of Cambridge’s Animal Welfare & Ethical Review Body Standing Committee. Experiments were performed under Home Office licenses (PPL8002474 and PPL8156839). All mice were bred as heterozygous crosses, and both male and female mice were used in experiments. Dat^Cre^ Fbxo^+/-^ and Dat^Cre^ Fbxo7^fl/-^ mice were generated as previously described (4).

### Plasmids

pAAV-CAG-Flex-mRuby2-GSG-P2A-GCaMP6s-WPRE-pA was a gift from Tobias Bonhoeffer & Mark Huebener & Tobias Rose (Addgene plasmid # 68717; http://n2t.net/addgene:68717; RRID:Addgene_68717). This construct was modified to remove GCaMP6s and replace it with mFbxo7. To achieve this, the MmFbxo7 gene (Image clone # 5054087) open reading frame was cloned in-frame with mRuby and P2A to generate an mRuby-P2A-Fbxo7 sequence. This sequence was then inserted in an antisense orientation relative to the chicken β-actin promoter and flanked by LoxP sites on both sides. Upon Cre recombinase expression, the transgene is flipped to the sense orientation, resulting in the activation of Fbxo7 and mRuby expression.

### rAAV Production

The viral particles were produced by Penn Vector Core (Gene Therapy Program Perelman School of Medicine, University of Pennsylvania, USA). To generate the Control Empty Vector and Fbxo7 viruses, we provided Penn Vector Core with the AAV9-PHP.eB.CAG.DIO.P2A.mRuby (Empty Vector, EV Virus) and AAV9-PHP.eB.CAG.DIO.FBXO7.P2A.mRuby (Fbxo7 Virus). The plasmid DNA was purified using an EndoFree Midi Prep Kit (Qiagen, Cat#12362) to ensure low endotoxin levels, suitable for viral production. The titer of the viral preparations obtained were 5.50x10^13^ genome copies (GC)/mL for the Empty Vector virus (EV) and 7.52x10^12^ GC/mL for the Fbxo7 virus.

### rAAV Injections

For tail vein injection, 4-or 9-week-old *Dat* ^*Cre*^ *Fbxo7* ^−/fl^ (KO) mice were injected with 7.52x10^11^ to 5.5x10^12^ GC of the empty virus (EV) or the Fbxo7 virus, in a total volume of 180 μL phosphate buffer saline. After the injection, mice were left to recover for 6 weeks before extraction of the brain for analysis.

### Tissue processing

Brains were harvested at the age of 10 weeks or 15 weeks, as indicated. Mice were euthanized with a 0.3 mL intraperitoneal injection of pentobarbitone sodium, 200 mg/ml (Euthatal, Merial, UK). They were perfused trans-cardially (10) with phosphate buffer saline followed by ice-cold 4% paraformaldehyde in phosphate buffer saline, pH 7.4. Brains were removed and post-fixed in 4% PFA overnight before being switched to 30% sucrose.

### Immunohistochemistry

Perfused brains were sectioned coronally at 35 µm intervals using a sledge microtome (Leica, CM 1950). Sections were collected in 12 series and washed in 0.1 M phosphate buffer saline before being incubated with primary and secondary antibodies (in 5% serum/0.05% Triton). Antibodies used were rabbit anti-Tyrosine Hydroxylase (TH) (Merk, ab152, 1:400) and Alexa Fluor 488 (Invitrogen, 1:400). Sections were visualized by diaminobenzidine (DAB; Cell Signaling Cat# 8059) according to the manufacturer’s protocol) or immunofluorescence (Alexa Fluor 488).

### Optical density analysis

To determine the fibre density in the striatum, the mean optical intensity was measured from the TH-positive stained sections. All DAB-stained slides were imaged using the Nanozoomer-XR slide scanner (Hamamatsu Photonics, Japan), at a resolution of 0.23um/pixel using a 40X objective. The areas of interest were the dorsal striatum and the nucleus accumbens. For the striatum, the measurements were made on 12 coronal sections, delineated as being between +1.7 and -1.4 relative to bregma. The lining of the ventricular wall and external capsule represented the medial and dorsal/lateral borders of the defined area of interest in the striatum. The ventral limit of the striatum was a diagonal line passing above the anterior commissure, between the external capsule and the ventricular wall. For more posterior sections, the lateral aspect of the globus pallidus was used as the medial border of the striatum, and a horizontal line was made between the external capsule and the globus pallidus to outline the ventral border. Density measurements for the nucleus accumbens were made across 4 coronal sections, approximately +1.7 to +1.0, relative to bregma. A diagonal line passing down through the anterior commissure, between the external capsule and the lowest portion of the ventricular wall, was used as the dorsal border and the lowest limits of the olfactory tubercle defined the ventral perimeter of the area we defined as the nucleus accumbens. TH staining along the lateral stripe of the striatum represented the lateral wall, and TH+ shell of the nucleus accumbens was used as the medial limits. All the fibre density analysis was done using ImageJ software.

### Statistical analysis

Statistical analysis was performed using GraphPad Prism software. All comparisons between groups of mice were performed using a One-way Anova, Tukey test. The number of animals used for each test is indicated in the results section. Differences were considered significant when the P vales were <0.05 (*p<0.05; **p<0.005, ***p<0.0005). Error bars on the graphs represent the standard deviation.

## Acknowledgements

We thank members of the Laman Lab for helpful discussions and dedicate this work, which was so adversely affected by the SARS-CoV-2 pandemic, to the little ray of hope who came into the world at that time. This work is supported by funding from Parkinson’s UK grant G-1701 and the Rosetrees Trust to HL and RAB. This research is also supported by the NIHR Cambridge Biomedical Research Centre (BRC-1215-20014, including the Cell Phenotyping Hub. The views expressed are those of the author(s) and not necessarily those of the NIHR or the Department of Health and Social Care. For the purpose of Open Access, the author has applied for a CC BY public copyright license to any Author Accepted Manuscript version arising from this submission.

